# How Fishing Cats *Prionailurus viverrinus* fish: Describing a felid’s strategy to hunt aquatic prey

**DOI:** 10.1101/2020.04.24.058925

**Authors:** Divyajyoti Ganguly, Tiasa Adhya

**Affiliations:** Department of Zoology, Maulana Azad College, Calcutta University, 8, Rafi Ahmed Kidwai Rd, Taltala, Kolkata, West Bengal 700013(DG); Centre for Conservation of Natural Resources, The University of Trans-disciplinary Health Sciences and Technology, 74/2, Post Attur via Yelahanka, Jarakabande Kaval, Bengaluru, Karnataka 560064 (TA); The Fishing Cat Project, Small Wild Cat Conservation Foundation, 90 Rincon Rd, Corrales, NM,87048-7619USA(TA)

**Keywords:** camera-trap, ethogram, felids, Fishing Cat, hunting, participatory-science

## Abstract

Hunting strategies are key to carnivore survival (Krebs and Davies 2009;Kamil et al. 2012;Michalko and Pekar 2016). Fishing Cat’s *(Prionailurus viverrinus)* persistence in the ‘semi-aquatic niche’ (Kitchener et al. 2010) despite felids being terrestrial carnivores in general (>95%) (Hunter 2019) suggests the evolution of a successful hunting strategy. Its further suggest selection for hunting in wetlands. What energy conserving strategies did the Fishing Cat borrow from its family and how were these adapted to optimize energy gained from hunting fish, its primary prey? We attempt to answer this by analyzing 197 video footages collected opportunistically from a participatory science initiative conducted over 2.5 years. We found that the felid switches between stationary and active modes of searching for prey depending on the depth of water and the corresponding loss of body heat/energy. For example, diving in deeper waters requires the submergence of the upper portions of the body and loss of more body heat/energy. Our analysis shows that the cat spent much of its time (~52%) sitting and waiting for prey (fish) to come nearer and then took limited attempts to dive into deeper water (2.78%). We suggest that this is a strategy to optimize the net energy gain. In shallow waters where the cat could forage without submerging the upper body it adopted a predominantly active mode of hunting (~96%) to flush out prey. Thus, prominent hunting strategies in the small cat lineage like ‘sit-and-wait’ and ‘active foraging’ is adapted to hunt in the water. We recorded a 60% hunting success in deeper waters but did not detect a successful hunt in shallow waters due to the low sample size of data from shallow water hunting. The major caveat in our study is the post-hoc analysis of opportunistically collected data as opposed to data derived from a structured design with predefined objectives. With more sampling from various seasons and landscapes, finer details can be explored which would have conservation implications. For example, we would expect variations in ‘attempts to hunt’ during cold seasons because heat loss might be less of a challenge in the latter. Quantifying ‘attempts to hunt’ and ‘successful hunts’ across seasons could help focus management interventions to minimize negative interactions between fish cultivators and Fishing Cat. The strictly nocturnal activity of fishing cat as demonstrated in our study could be a strategy taken by the cat to avoid humans. Our approach of using participatory-science is relevant for conducting research on mammal behavior in human dominated landscapes.

---

Successful hunting strategies allow the persistence of efficient predators through natural selection (Krebs and Davies 2009;Kamil et al. 2012;Michalko and Pekar 2016). The guild exhibits a diversity of such hunting strategies that optimize prey capture while overcoming foraging constraints (Hayward et al. 2006;Carbone et al. 2007;Williams et al. 2014;Rizzuto et al. 2018). For example, some members of Canidae hunt in groups to prey by coordinating a prolonged chase in open areas (Kruuk and Turner 1967;Murray. 1995;Padilla and Hilton 2015). In contrast, Felidae mostly has solitary hunters that ‘stalk’ distance to prey before a successful ambush (Carbone et al. 2007;Sunquist and Sunquist 2013). They do this with the help of cover in their immediate surroundings (Kruuk and Turner 1967;Elliot et al. 1977;Corbett 1979;Thiel 2011). With the exceptions of Cheetah (*Acinonyx jubatus*) (Sunquist and Sunquist 2013) and Snow Leopard (*Panthera uncia*) (Sunquist and Sunquist 2017), felids do not engage in long chases.

A brief chase is present in some large felids while others hunt by pouncing or with a short rush (Kruuk and Turner 1967;Sunquist and Sunquist 2017). Large felids in general hunt prey their body size (Carbone et al. 2007;Sunquist and Sunquist 2017). Chase is still less small felid (<25 kg) hunting strategies (Table 1) including in the larger and heavier ones (15-25 kg) that can opportunistically take prey similar to their body size (Carbone et al.2007; Sunquist and Sunquist 2017;Hunter 2019). Their diet is primarily composed of prey such as rodents, lagomorphs, birds, reptiles and invertebrates (Sunquist and Sunquist 2017;Hunter 2019) that require lesser handling time, i.e., less energy (Carbone et al. 2007). Hence, capturing prey through short bursts of chase or by concentrating efforts in a to hunt such as through leap/dash/pounce is common (Table 1). This counters the greater amount of energy spent in searching and procuring smaller prey by small carnivores (Carbone. 2007;Rizzuto et al. 2018). In addition, they lose more body heat (energy) due to a area/body mass ratio (Tracy 1977).

**Table 1.**
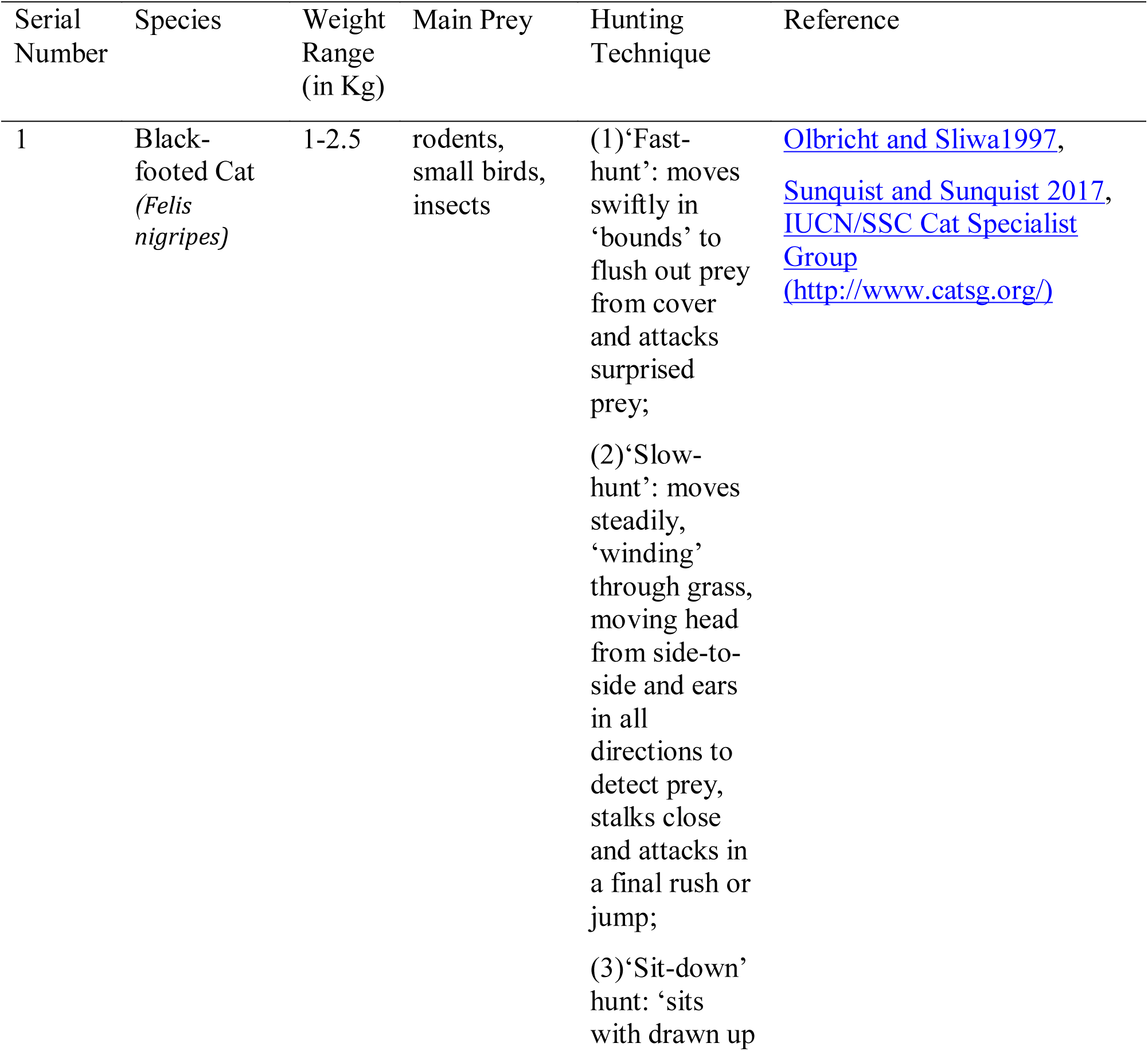
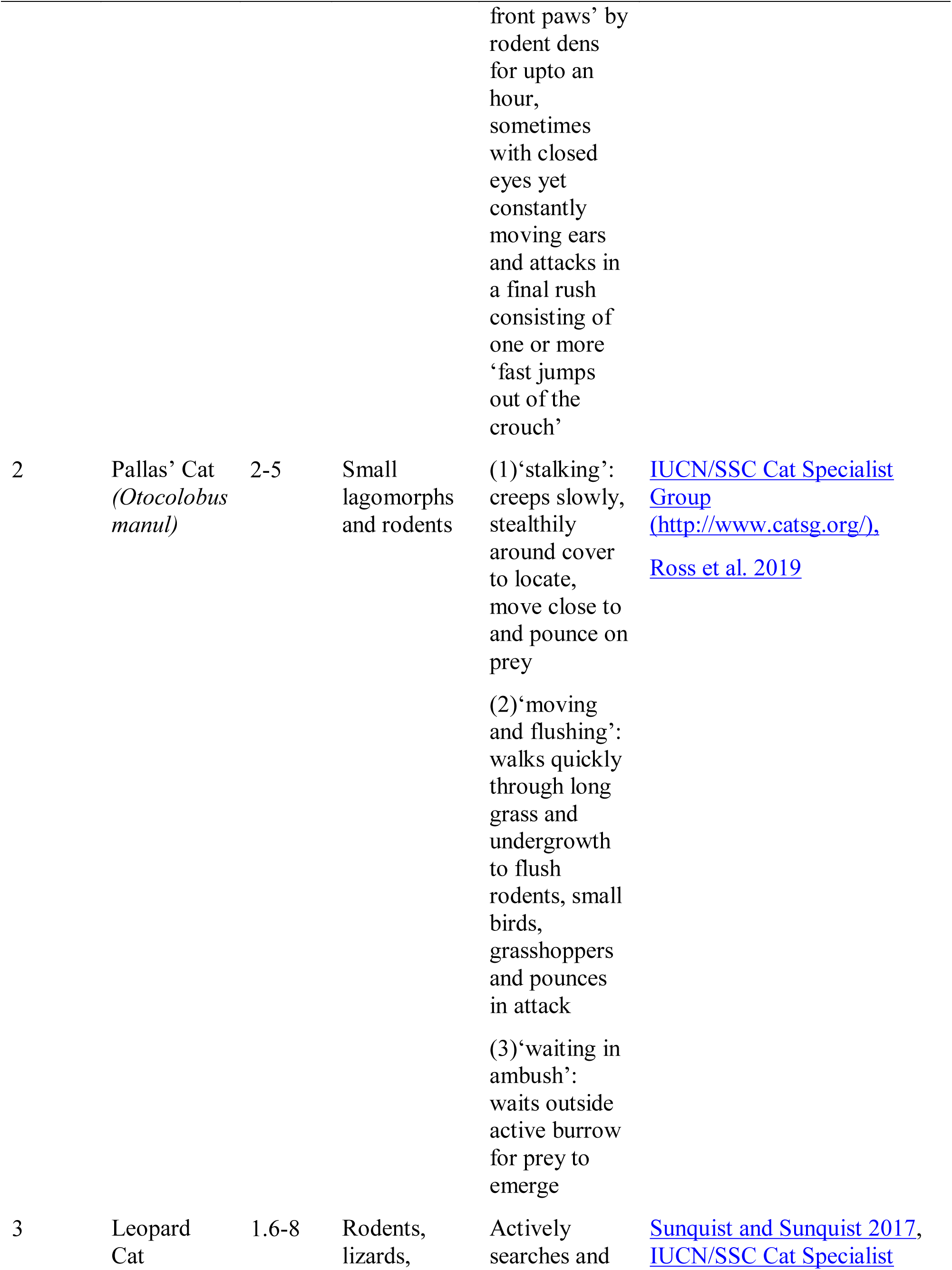
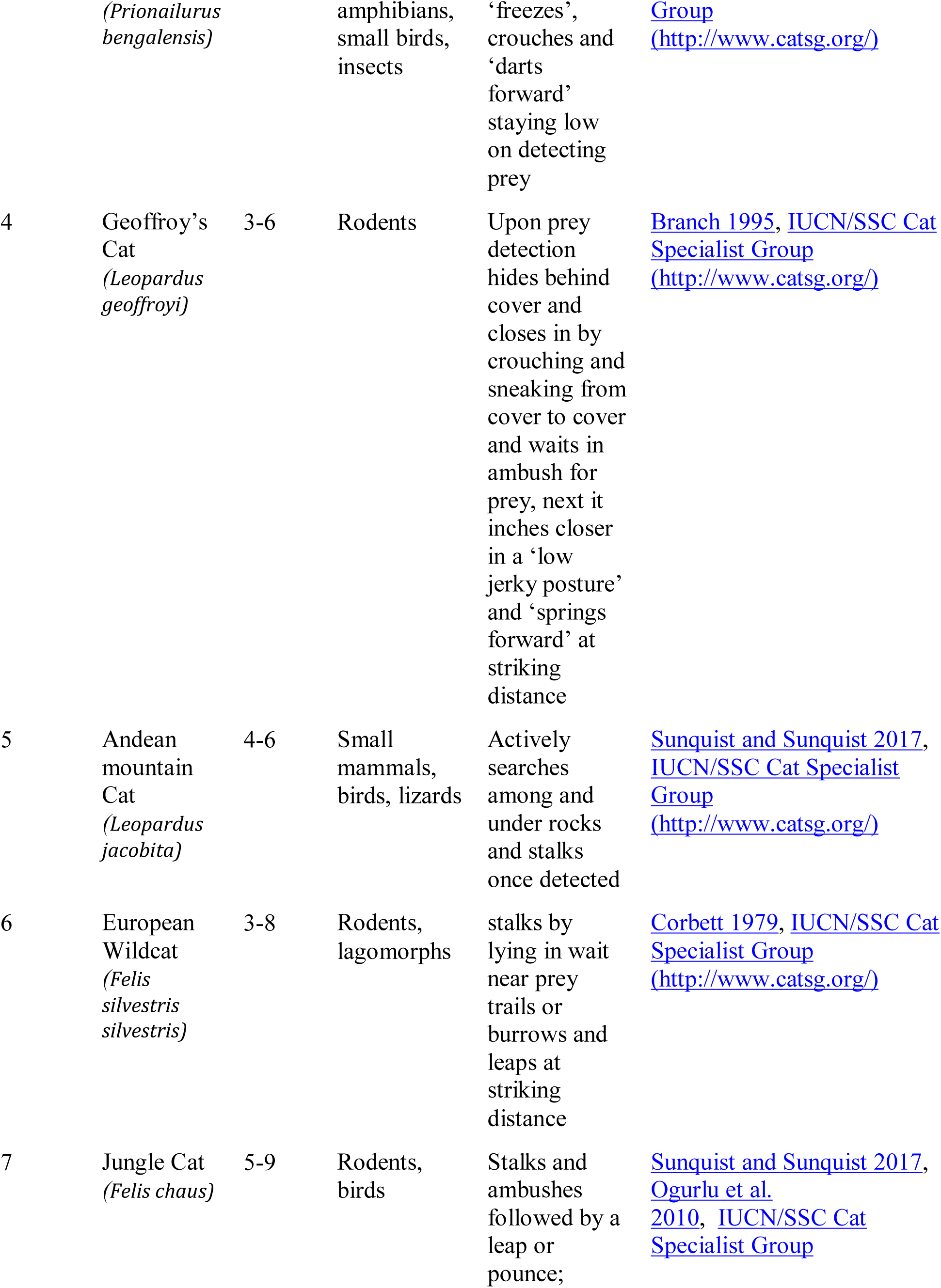
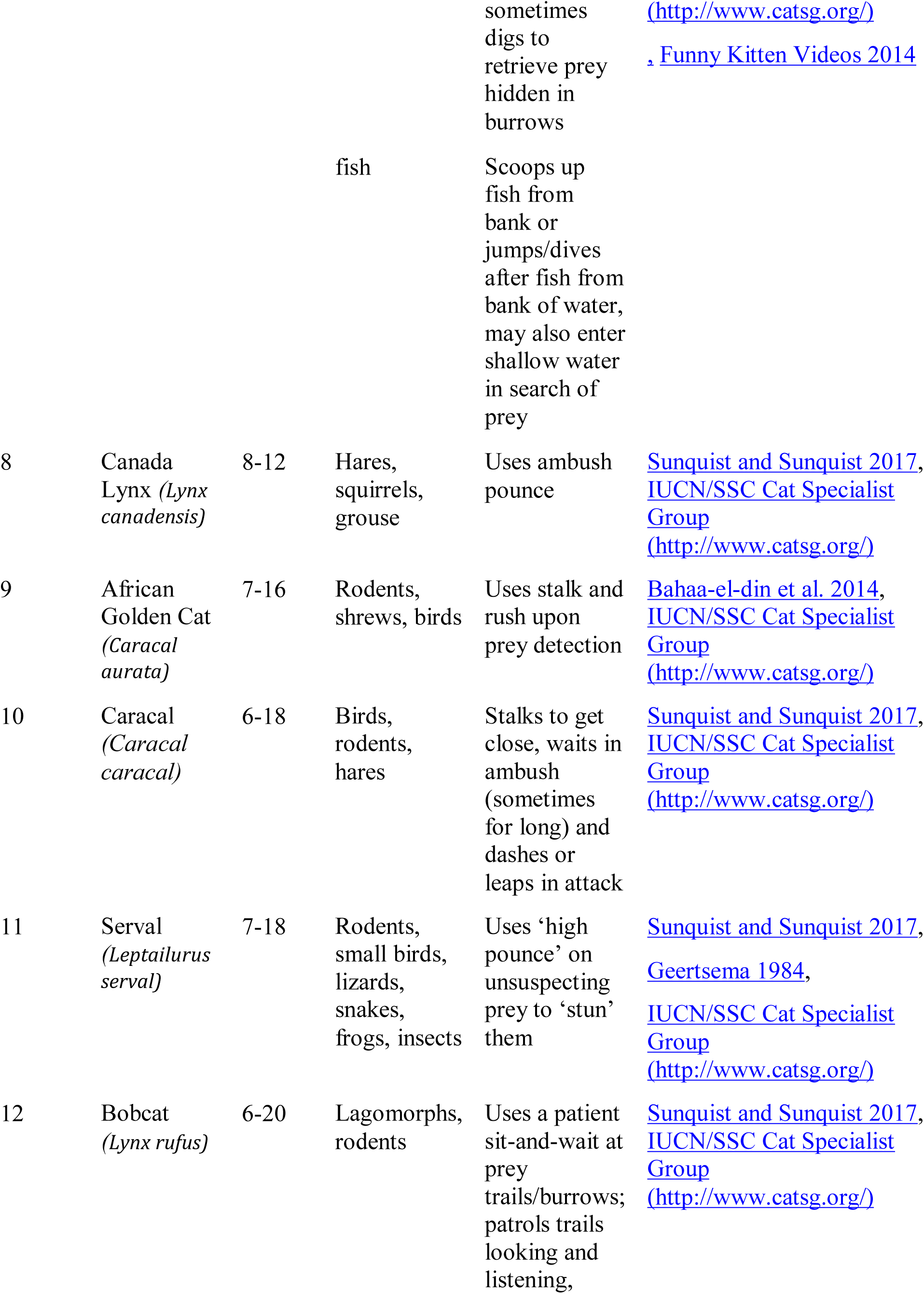
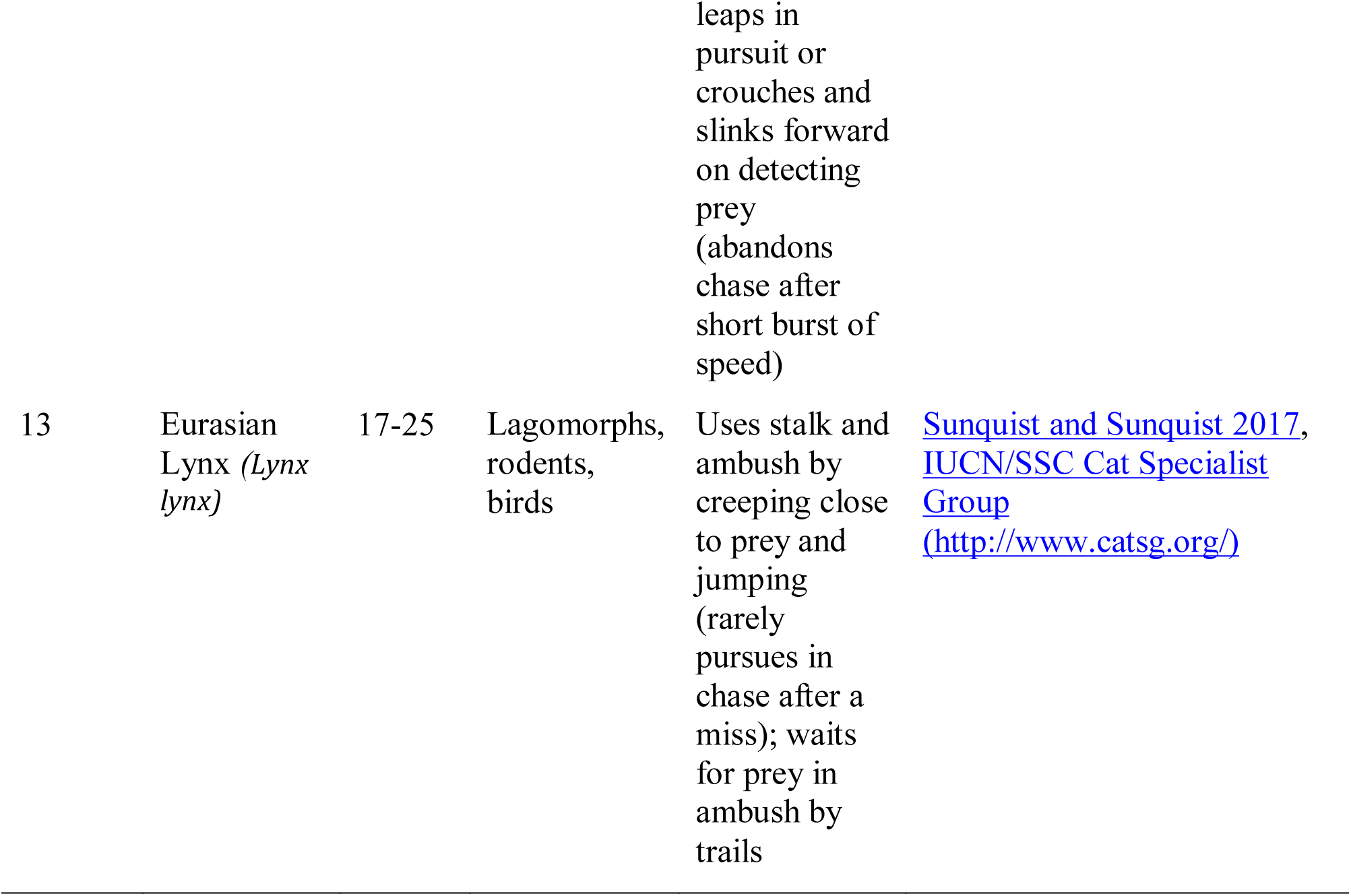
Weight, prey size and hunting technique in small felids

| Serial Number | Species | Weight Range (in Kg) | Main Prey | Hunting Technique | Reference |
| --- | --- | --- | --- | --- | --- |
| 1 | Black-footed Cat (*Felis nigripes*) | 1-2.5 | rodents, small birds, insects | (1)‘Fast-hunt’: moves swiftly in ‘bounds’ to flush out prey from cover and attacks surprised prey;<br>(2)‘Slow-hunt’: moves steadily, ‘winding’ through grass, moving head from side-to-side and ears in all directions to detect prey, stalks close and attacks in a final rush or jump;<br>(3)‘Sit-down’ hunt: ‘sits with drawn up front paws' by<br>rodent dens<br>for upto an<br>hour,<br>sometimes<br>with closed<br>eyes yet<br>constantly<br>moving ears<br>and attacks in<br>a final rush<br>consisting of<br>one or more<br>'fast jumps<br>out of the<br>crouch' | Olbricht and Sliwa1997,<br>Sunquist and Sunquist 2017,<br>IUCN/SSC Cat Specialist Group<br>[(http://www.catsg.org/)](http://www.catsg.org/) |
| 2 | Pallas' Cat<br>(*Otocolobus*<br>*manul*) | 2-5 | Small<br>lagomorphs<br>and rodents | (1)'stalking':<br>creeps slowly,<br>stealthily<br>around cover<br>to locate,<br>move close to<br>and pounce on<br>prey<br>(2)'moving<br>and flushing':<br>walks quickly<br>through long<br>grass and<br>undergrowth<br>to flush<br>rodents, small<br>birds,<br>grasshoppers<br>and pounces<br>in attack<br>(3)'waiting in<br>ambush':<br>waits outside<br>active burrow<br>for prey to<br>emerge | IUCN/SSC Cat Specialist<br>Group<br>[(http://www.catsg.org/)](http://www.catsg.org/),<br>Ross et al. 2019 |
| 3 | Leopard<br>Cat | 1.6-8 | Rodents,<br>lizards, | Actively<br>searches and | Sunquist and Sunquist 2017,<br>IUCN/SSC Cat Specialist |
|  | *(Prionailurus bengalensis)* |  | amphibians, small birds, insects | ‘freezes’, crouches and ‘darts forward’ staying low on detecting prey | [Group](http://www.catsg.org/) |
| 4 | Geoffroy’s Cat<br>*(Leopardus geoffroyi)* | 3-6 | Rodents | Upon prey detection hides behind cover and closes in by crouching and sneaking from cover to cover and waits in ambush for prey, next it inches closer in a ‘low jerky posture’ and ‘springs forward’ at striking distance | [Branch 1995, IUCN/SSC Cat Specialist Group](http://www.catsg.org/) |
| 5 | Andean mountain Cat<br>*(Leopardus jacobita)* | 4-6 | Small mammals, birds, lizards | Actively searches among and under rocks and stalks once detected | [Sunquist and Sunquist 2017, IUCN/SSC Cat Specialist Group](http://www.catsg.org/) |
| 6 | European Wildcat<br>*(Felis silvestris silvestris)* | 3-8 | Rodents, lagomorphs | stalks by lying in wait near prey trails or burrows and leaps at striking distance | [Corbett 1979, IUCN/SSC Cat Specialist Group](http://www.catsg.org/) |
| 7 | Jungle Cat<br>*(Felis chaus)* | 5-9 | Rodents, birds | Stalks and ambushes followed by a leap or pounce; | [Sunquist and Sunquist 2017, Ogurlu et al. 2010, IUCN/SSC Cat Specialist Group](http://www.catsg.org/) |
|  |  |  |  | sometimes<br>digs to<br>retrieve prey<br>hidden in<br>burrows | [(http://www.catsg.org/)](http://www.catsg.org/)<br>. Funny Kitten Videos 2014 |
|  |  |  | fish | Scoops up<br>fish from<br>bank or<br>jumps/dives<br>after fish from<br>bank of water,<br>may also enter<br>shallow water<br>in search of<br>prey |  |
| 8 | Canada<br>Lynx (*Lynx*<br>*canadensis*) | 8-12 | Hares,<br>squirrels,<br>grouse | Uses ambush<br>pounce | Sunquist and Sunquist 2017,<br>IUCN/SSC Cat Specialist<br>Group<br>[(http://www.catsg.org/)](http://www.catsg.org/) |
| 9 | African<br>Golden Cat<br>(*Caracal*<br>*aurata*) | 7-16 | Rodents,<br>shrews, birds | Uses stalk and<br>rush upon<br>prey detection | Bahaa-el-din et al. 2014,<br>IUCN/SSC Cat Specialist<br>Group<br>[(http://www.catsg.org/)](http://www.catsg.org/) |
| 10 | Caracal<br>(*Caracal*<br>*caracal*) | 6-18 | Birds,<br>rodents,<br>hares | Stalks to get<br>close, waits in<br>ambush<br>(sometimes<br>for long) and<br>dashes or<br>leaps in attack | Sunquist and Sunquist 2017,<br>IUCN/SSC Cat Specialist<br>Group<br>[(http://www.catsg.org/)](http://www.catsg.org/) |
| 11 | Serval<br>(*Leptailurus*<br>*serval*) | 7-18 | Rodents,<br>small birds,<br>lizards,<br>snakes,<br>frogs, insects | Uses 'high<br>pounce' on<br>unsuspecting<br>prey to 'stun'<br>them | Sunquist and Sunquist 2017,<br>Geertsema 1984,<br>IUCN/SSC Cat Specialist<br>Group<br>[(http://www.catsg.org/)](http://www.catsg.org/) |
| 12 | Bobcat<br>(*Lynx rufus*) | 6-20 | Lagomorphs,<br>rodents | Uses a patient<br>sit-and-wait at<br>prey<br>trails/burrows;<br>patrols trails<br>looking and<br>listening, | Sunquist and Sunquist 2017,<br>IUCN/SSC Cat Specialist<br>Group<br>[(http://www.catsg.org/)](http://www.catsg.org/) |
| 13 | Eurasian<br>Lynx (*Lynx*<br>*lynx*) | 17-25 | Lagomorphs,<br>rodents,<br>birds | Uses stalk and<br>ambush by<br>creeping close<br>to prey and<br>jumping<br>(rarely<br>pursues in<br>chase after a<br>miss); waits<br>for prey in<br>ambush by<br>trails | Sunquist and Sunquist 2017,<br>IUCN/SSC Cat Specialist<br>Group<br>[(http://www.catsg.org/)](http://www.catsg.org/) |

Small felids have evolved different hunting strategies depending on the prey taken. For instance, small mammals like rodents and hares are ambushed by sitting and waiting near burrows or along trails frequented by prey or even by digging at their burrows (Sunquist and Sunquist 2017). Birds on the ground are either flushed out from vegetation or stalked upon distance (Olbricht and Sliwa 1997;Sunquist and Sunquist 2017). Arboreal primates by climbing on trees and scaring them to lower branches and then to the ground as seen in Ocelot (*Leopardus pardalis*) (Bianchi and Mendes 2007) whereas Margay (*Leopardus wiedii*) uses vocal mimicry to lure prey (de Oliveira Calleia and Gordo 2009). Kitchener et al. (2010) sequentially divide a typical felid hunt into ambush, detecting prey, stalking and running, catching, killing, processing and eating. While ambush, detecting prey and stalking are typically low-cost activities, running and catching are associated with higher costs (Williams et al. 2014). The cost of killing, processing and eating differs with the type and size of prey. Small rodents or birds can be easily subdued and consumed whole while large prey like large birds are required to be processed by removal of feathers before eating (Sunquist and Sunquist 2017).

Fishing Cat (*Prionailurus viverrinus*) and Flat-headed Cat (*Prionailurus planiceps*) are exceptional because of their choice of aquatic prey (Kitchener et al. 2010) as against taken by most felids. Further, morphological adaptations such as semi-retractile claws to grip slippery aquatic prey and a double-coated fur to prevent the body from getting wet suggest strong selection for hunting in wetlands (Hunter 2019). However, dentition in Fishing C a more eclectic diet when compared to Flat-headed Cat’s (Kitchener et al. 2010;Hunter 2019). The muscular build of Fishing Cat is suitable for hunting down similar sized (Sunquistand Sunquist 2017;Hunter 2019). Despite a broad dietary spectrum, fish is dietary component (Haque and Vijayan 1993;Cutter 2015;Hunter 2019). Examining strategy of the cat could provide insight on why such a preference could have evolved. The elusive nature of the cat however is an impediment for studying their behavior in the wild (Sunquist and Sunquist 2017). Only a few observations on their hunting exist (Prater 1965;Sunquist and Sunquist 2017;Hunter 2019). Technological advances such as camera traps have however facilitated non-invasive studies on their ecology (Nair 2012;Das et al. 2017;Malla 2016;Malla et al. 2019). But it is difficult to conduct such studies dominated landscapes due to the issues of theft and requirement of greater manpower to prevent the same.

Here, we used data collected through a participatory-science initiative. Such initiatives are targeted at achieving answers to real world questions under limited resources, in a cost-effective way through a partnership between volunteers and scientists where volunteers contribute data and/or assist with its analysis (Miller-Rushing et al. 2012;Lawson et al. 2015). Such initiatives find use in a wide variety of wildlife disciplines. For example, ornithology has benefited the most from citizen science initiatives with data on relative abundance and distribution of bird species (Lawson et al. 2015). Participatory-science has also found use on wildlife management (van Vliet et al. 2018), land-use change (Johansson and Isgren 2017), wildlife monitoring (de Mattos Vieira et al. 2015), and wildlife disease surveillance (Lawson et al. 2015).

We engaged residents sharing space with Fishing Cat in documenting and monitoring the cats in their backyard through a camera trap exercise (http://thelastwilderness.org/know-thy-neighbours-the-spectacular-fishing-cat/). We used opportunistically collected camera-trap data 2.5 years from this program along with field observations. All the behavior types from the Fishing Cat footages were first identified and used in ethogram construction. An ethogram is a formal description in the form of a table of all behavior types observed in a species or of particular functional classes of behavior(McDonnell and Poulin 2002).

The major objectives of our study were to:

a. describe major behavioral activities of Fishing Cat around waterbodies and channels through ethogram
b. examine activity budget in order to describe the hunting strategy of Fishing Cat

## Materials and Methods

Study area.—Our study area consists of 3 sites, the Howrah district in the lower Gangetic floodplains of West Bengal and two coastal wetlands in the Mahanadi floodplains along the Eastern coast - Paradip and Chilika (Fig. 1).

**Fig. 1.**
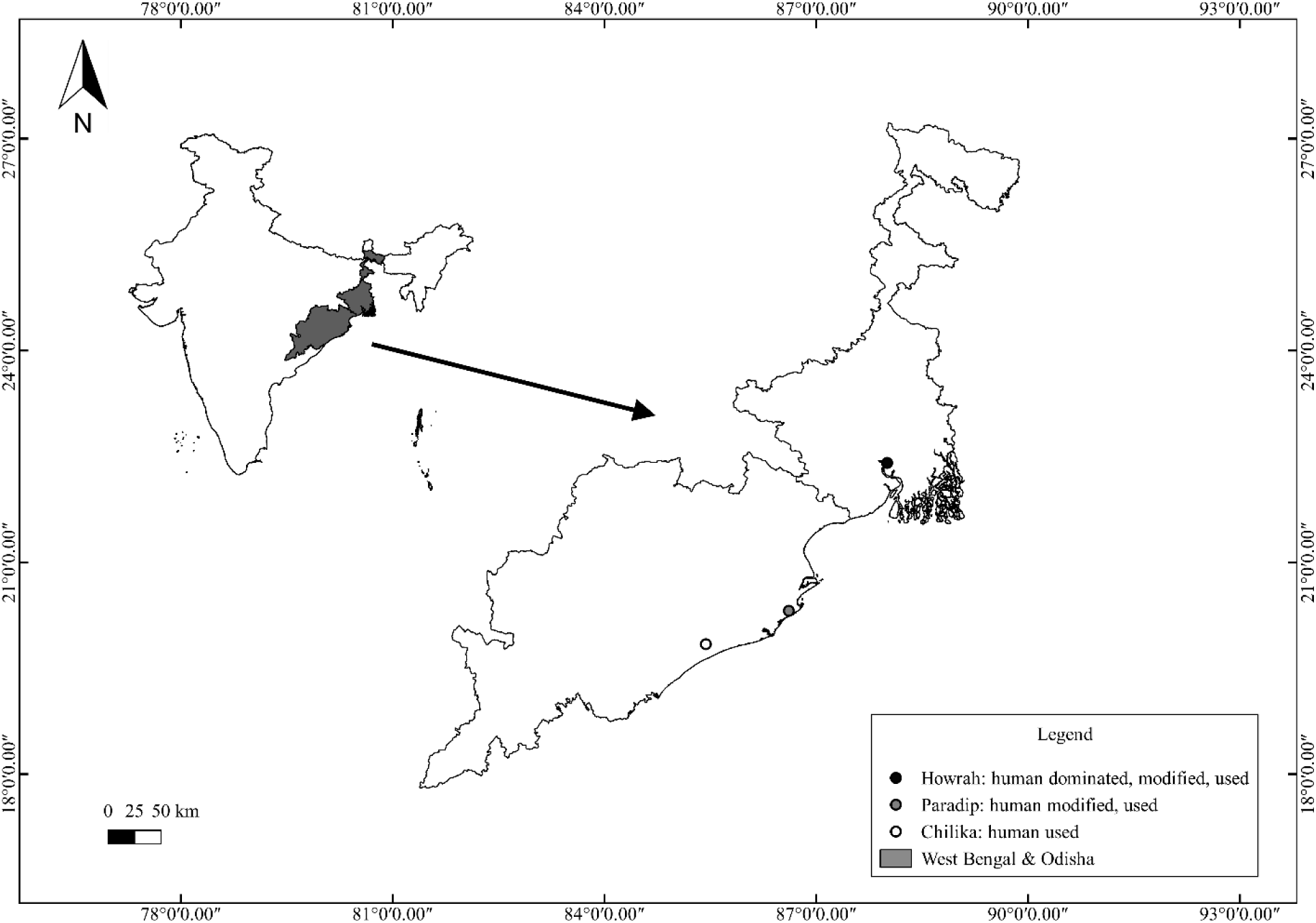
Map showing study area sites (circled) across West Bengal and Odisha. Howrah, Paradip and Chilika are marked with Black, grey and white circles respectively

The Howrah district (22°27’40.66“N, 88° 0’32.71”E) lies in the lower Gangetic floodplains and is drained by the Hooghly and the Rupnarayan rivers. Our study was conducted in fragmented marshy patches of Howrah district interspersed with croplands, small scale aquaculture, human settlements and industrial complexes. Human density of Howrah district is also highest among all sites and due to the sub-urban nature of the populace, human activity is high till 10 pm.

In Paradip (20°18’50.63“N, 86°36’45.49”E) data was collected from a marshy area of 9 sq. km inside the Paradeep Phosphates Limited campus. Small-scale aquaculture ponds intersperse the marshes. Fishing activity by humans take place during day-time. After dark, human activity is minimized since the area is accessible to campus residents only.

The study site in Chilika (19°51’39.66“N, 85°25’32.95”E) located along the lake’s north east sector constitutes coastal wetlands formed by the Mahanadi floodplains in the Eastern coast and has a contiguous marshland of 115 sq. km., crisscrossed by canals. Here local fishermen catch wild fish during the day or anchor their boat in water channels at night after laying nets and/or fish-baskets. The area is not inhabited by humans and all human activity stops save for travelling fishing boats early morning. Human activity and presence is minimal in Chilika when compared to Howrah and Paradip.

*Data collection*.—We initiated a participatory-science initiative in the study sites among interested and enthusiastic members of the local community who were given camera monitor Fishing Cat in their neighborhood. The initiative was named ‘Know Thy Neighbour’. 20 camera traps (14 Cover Illuminator 9 and 6 Browning IR) were deployed opportunistically over 2.5 years starting from November 2016 across the different study sites. Camera traps were deployed to maximize the probability of capturing Fishing Cat. We consulted local people to decide where to deploy the camera traps. As suggested by them we searched for probable sites to place camera traps along shallow ponds or water channels. All camera trap deployment sites were chosen after locating scats and/or tracks of felids. They were set on video mode and were active from 5:30 p.m. - 6:00 a.m. Each trained volunteer picked up traps early morning and repositioned them at the same location at night to reduce chances of misplacement/theft. We assumed that we would not fail to miss fishing cat activity when camera traps were non-active since Fishing Cats are primarily nocturnal (Hunter 2019) and villagers mostly reported after dark.

*Data analysis*.—We collected over 200 camera trap video footages from the participatory science initiative to analyze hunting behavior of Fishing Cat. All videos were analyzed by the first author to avoid introducing observer bias. Consecutive videos from the same camera trap on the same night which were taken less than a minute apart were considered as single data, as we assumed lack of independence between closely occurring activities. ‘Behavior States’ were first identified which represents behavioral patterns of relatively long duration (Martin and Bateson 1993). These were further divided into ‘behavior sub-states’. Behavioral patterns of duration were classified as Events (Martin and Bateson 1993). We calculated of time spent performing each State and the frequency of each Event using an ethogram.

To examine activity pattern, videos (n=98) were selected such that they differed by at least 30 minutes to ensure independence between data points. Such videos were then used to generate line graphs to understand peak activity pattern of Fishing Cat. If two separate individuals were detected in the videos, these were considered independent behavioural events and analysed as two different samples.

In addition to these, any visible social behavior involving interactions between two or more individuals was also noted.

In total, we analyzed 139.92 minutes from 197 camera trap video footages constituting 198 datapoints from all three study sites combined.

## RESULTS

This resulted in the identification of 12 distinct ‘Behavior States’ with their components, i.e., behavior sub-types. The 12 ‘Behavior States’ are ‘Stationary Searching’ (with behavior sub-types - ‘Standing’, ‘Sitting’, ‘Crouching’), ‘Mobile Searching’ (behavior sub-types: ‘Slow Walking’, ‘Fast Walking’), ‘Approaching’, ‘Stalking’ (behavior sub-types: ‘Stationary Stalking’, ‘Mobile Stalking’), ‘Attempt to hunt/Attacking’, ‘Eating Prey’, ‘Pheromone Spray’, ‘Defecating’, ‘Sniffing’, ‘Self-Grooming’, ‘Swimming’ and ‘Yawning’. We found 7 States and 5 Events (Table 3 and 4). The detailed ethogram is presented in Table 2.

**Table 2.** Ethogram of Fishing Cat’s behaviour around resource patches (water body), definitions from Stanton et al. 2015

| Behavior | Description | Supplementary Data |
| --- | --- | --- |
| **Slow Walking** | Forward locomotion along water body at a normal gait with occasional scanning of water body | Supplementary Data SD 2 |
| **Fast Walking** | Forward locomotion along water body at a swift gait, faster than slow walking | Supplementary Data SD 1 |
| **Approaching** | Slow and forward locomotion, sometimes in a slightly crouched position, directed towards water body with eyes and ears moving and scanning the water | Supplementary Data SD 5 |
| **Standing** | All four legs extended and paws touching ground along or within water | Supplementary Data SD 3 |
| **Sitting** | Hind legs flexed and resting on the ground, front legs extended, body straight or lying on the ground against on side of body with head raised, along water | Supplementary Data SD 7 |
| **Crouching** | Body close to the ground, whereby all the legs remain bent and belly touching or slightly raised above the ground along water body | Supplementary Data SD 4 |
| **Mobile Stalking** | Slow and forward locomotion in a slightly crouched position, head low, eyes and ears focused on prey (movement in water) | Supplementary Data SD 6 |
| **Stationary Stalking** | Eyes and ears focused while stationary along water-edge | Supplementary Data SD 7 |
| **Attempt to hunt/Attacking** | Either jumping into water from sitting/standing position, or, charging towards and plunging front legs first into water | Supplementary Data SD 7 |
| **Eating prey** | Prey is ingested by chewing and swallowing | Supplementary Data SD 8 |
| **Pheromone spray** | Spray released backwards against vertical surface or object with vertically raised quivering tail while standing or in a squatting position on the ground, sometimes while walking along water body | Supplementary Data SD 7 |
| **Defecating** | Faecal matter deposited on the ground while in a squatting position after thoroughly sniffing ground along water body | Supplementary Data SD 12 |
| **Self-grooming** | Paw licked and wiped over head and body scratched using hind leg along water body | Supplementary Data SD 10 |
| **Sniffing** | Air inhaled through nose with movement of whiskers | Supplementary Data SD 4 |
| **Swimming** | Body propelled in water | Supplementary Data SD 11 |
| **Yawning** | Mouth widely opened while inhaling and closed while exhaling | Supplementary Data SD 13 |

**Table 3.** Proportion of time spent in performing each state as observed in deeper water, shallow water and combined by Fishing Cat

| State | Proportion of time spent (%)<br>{(Time spent in performing one state/Time spent in performing all states) X 100} |  |  |
| --- | --- | --- | --- |
|  | Deeper Water (n=180) | Shallow water (n=18) | Overall (n=198) |
| Mobile/Active Searching | 32.79 (Mobile) | 96.32 (Active) | 35.38 |
| Stationary Searching | 51.65 | — | 49.75 |
| Approaching | 5.15 | — | 4.73 |
| Stalking | 4.05 | 3.68 | 4.29 |
| Sniffing | 4.63 | — | 4.25 |
| Self-grooming | 1.38 | — | 1.25 |
| Swimming | 0.38 | — | 0.35 |

**Table 4.** Frequency of occurrence of each event as observed in Fishing Cat

| Event | Frequency of Occurrence |
| --- | --- |
| Attempt to hunt | 11 |
| Eating prey | 3 |
| Pheromone spray | 9 |
| Defecating | 1 |
| Yawning | 1 |

The felid spent much of its time (89.86%) searching for prey around resource patches (Fig. 2).

**Fig. 2.**
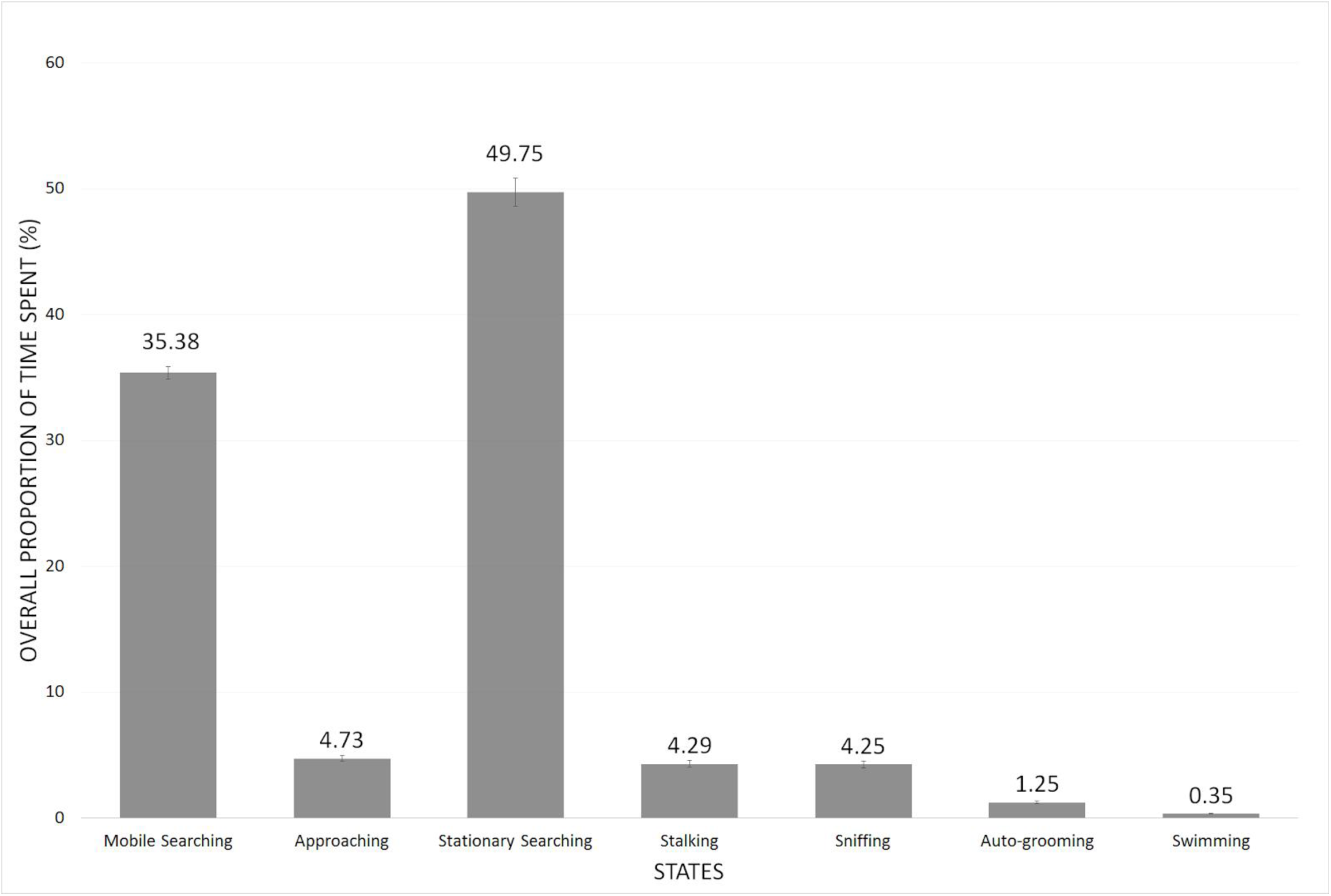
Proportion of time spent in performing each state (Mobile Searching, Approaching, Stationary Searching, Stalking, Sniffing, Self-Grooming, Swimming) by Fishing Cat, where error bars represent Standard Error (SE)

### Strategy in Deeper Water (n=180)

*Searching*.—Here it walked either fast or at a slower pace (‘Mobile Searching’ = 32.79%) along the banks of water bodies, stopping intermittently to observe the water surface and on a few occasions (n=9) to spray urine on bank vegetation. It also kept observing the edge of the water body, which was usually fringed with floating/submerged/emergent vegetation, from time to time while walking. On finding a suitable spot, it slowed down, sniffed the ground or air (4.63%) and approached (5.15%) closer towards the water body. ‘Approaching’ was sometimes followed by waiting (‘Stationary Searching’ = 51.65%). While waiting, which includes standing, sitting and crouching, the cat remained in ambush, keeping mostly still and carefully watching the water near the edge for signs of prey. Fishing Cat was also found to perch on beds of vegetation formed on banks presumably to catch fish entrapped in ‘dhauris’ or bamboo fish-baskets set by artisanal fishermen.

*Stalking.*—‘Approaching’ is occasionally followed by ‘stalking’ (4.05%). When prey is detected, it became alert and stalked with sight and ears focused in prey’s direction. While stalking, the cat either a) kept still and waited for prey to swim within striking distance (‘Stationary Stalking’), or, b) advanced closer on ground to reach within striking distance (‘Mobile Stalking’). In doing so floating vegetation might have provided some concealment to the cat.

*Attacking.*—At striking distance the fishing cat plunged/dived (depending on depth) into water (n=5) with forelimbs stretched followed by its head to catch prey. If the attack was unsuccessful, the cat immediately came out of water, shook off excess water from the body and in might take position immediately at a different spot.

### Strategy in Shallow Water (n=18)

*Searching.*—In knee-deep/paw-deep water the cat resorted to a more active form of search (‘Active Searching’, 96.32%) whereby it walked in the shallow water, pawing at pebbles or grass perhaps to flush out prey from hiding.

*Stalking.*—The cat stalked (3.68%) by keeping still for prey to come within striking distance, i.e., it only employed ‘Stationary Stalking’.

*Attacking.*—The cat pounced (n=6) after prey either post stalking or immediately on detecting movements while in ‘Active Searching’ mode. We did not record a successful attempt.

### Eating and Killing

In a successful attempt(n=3), the cat grabs hold of the prey and emerges out of water onto land. Small prey got consumed quickly and wholly along the bank while keeping vigilant.

However, large prey was dragged away from the water body as was observed in the field when a snake-head presumably around 2 kilograms in weight was caught by a female Fishing Cat and dragged into the nearby reeds. According to observations in the field, the cat consumed the flesh of larger fish while the head, midrib, intestines and egg-sac are left behind.

**Fig. 3.**
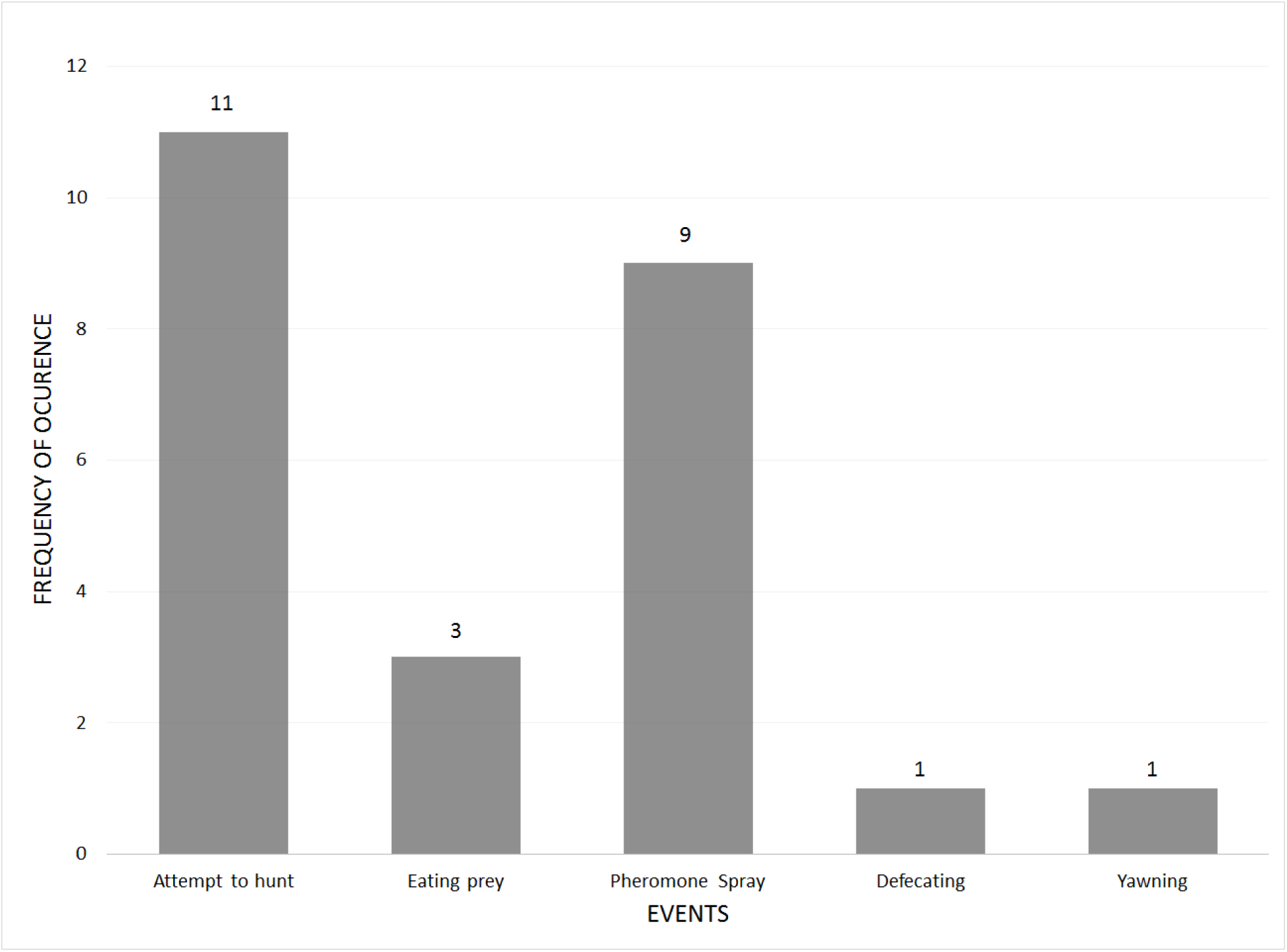
Frequency of occurrence of each state (Attempt to hunt, Eating prey, Pheromone Spray, Defecating, Yawning) by Fishing Cat

### Hunting Success

Overall, we recorded a naive hunting success rate of 27.27% (i.e., 3 successful attempts out of 11 events). Frequency of attempt to hunt were recorded directly from camera trap records and inferred from videos where the cat had fish-in-mouth. We assumed that an event of fish-in-mouth must have been preceded by a successful hunting attempt. ‘Attempt to hunt’ in deeper water was 2.78% and the same in shallow water was found to be 33.33%. Hunting success in deeper water was 60% (i.e., 3 successful hunts out of 5 attempts). No successful hunt was recorded from shallow water.

We found very little proportion of swimming behavior (0.35%).

Multiple individuals were found to use the same resource patch and were also found to patrol the same camera trap site in the same night. Individuals were also found to spray pheromones along vegetation, or, prominent structures like a stack of hay near the waterbody.

Cats engaged in ‘Self-Grooming’ sometimes (i.e., 1.25% of the time). Self-grooming is a maintenance behavior after feeding (Rochlitz 2014).

We also recorded instances of social interactions between fishing cat individuals. In one such instance, a male and a female were observed walking a trail, one behind another, along a pond (Supplementary Data SD 16). In another instance, two individuals were observed seemingly to communicate vocally by the side of a pond (Supplementary Data SD 14). A mother with two sub-adult kittens was also observed to interact in a series of video clips while foraging in shallow wetlands (Supplementary Data SD 15).

Nocturnal activity pattern of the fishing cat showed a gradual shift towards midnight with increasing levels of human presence in the area. In Howrah district, the area with most human presence and activity, Fishing Cat hunting times peak between 12 p.m. - 4 a.m. (higher peak; n=37; Fig. 4a). Paradip area with intermediate human presence, showed Fishing Cat activity peaking between 10-12 p.m. (n=4) and 2-4 a.m. (n=5) (Fig. 4b) and in Chilika area with the least amount of human presence, activity peaked between 6-8 p.m. (n=8) and 10-12 p.m. (n=7) (Fig. 4c). All three areas combined, Fishing Cat activity peaked between 12 p.m.-6 a.m. (n=59; Fig. 4d).

**Fig. 4a.**
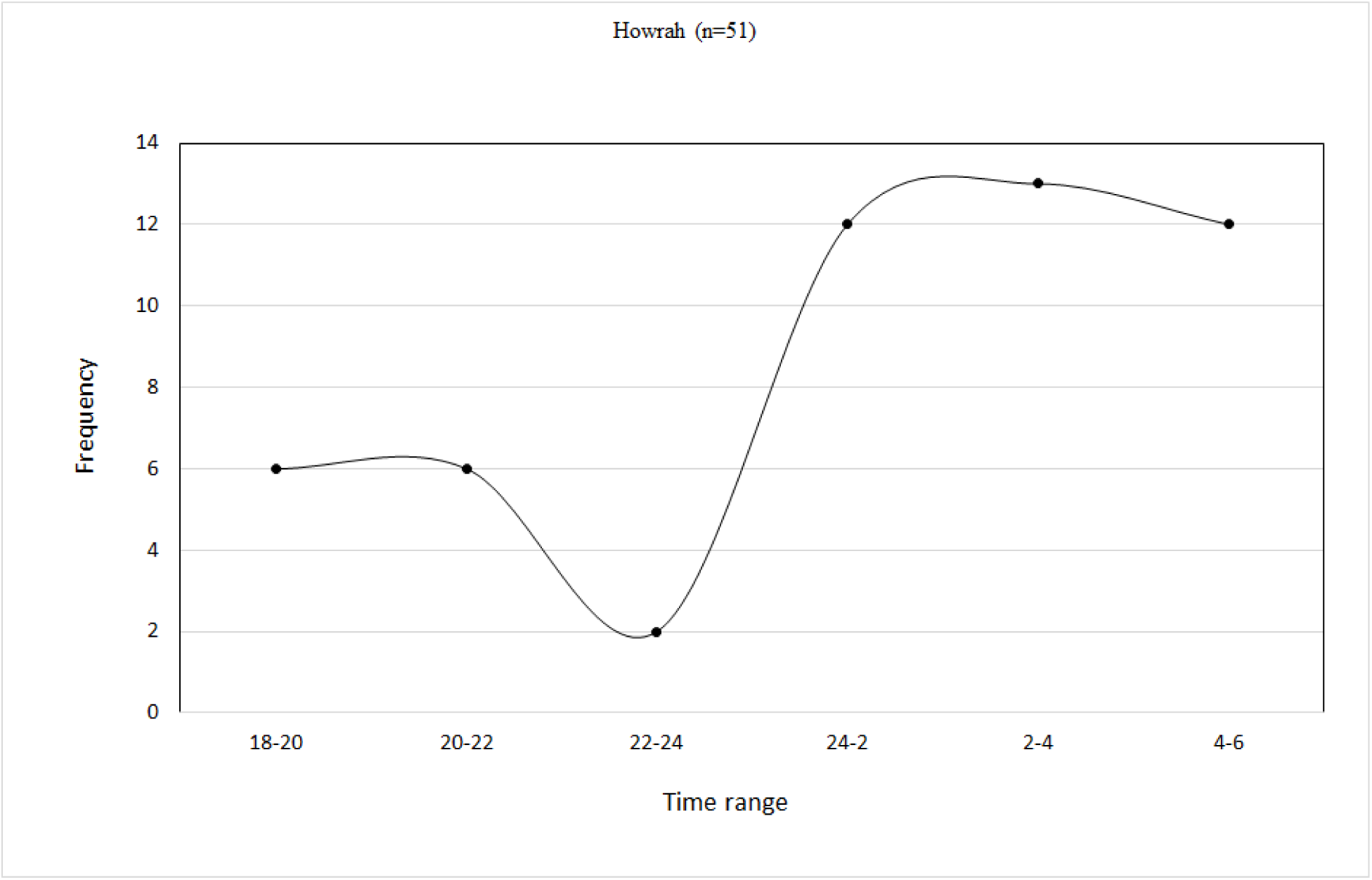
Graph showing activity pattern of Fishing Cat in Howrah

**Fig. 4b.**
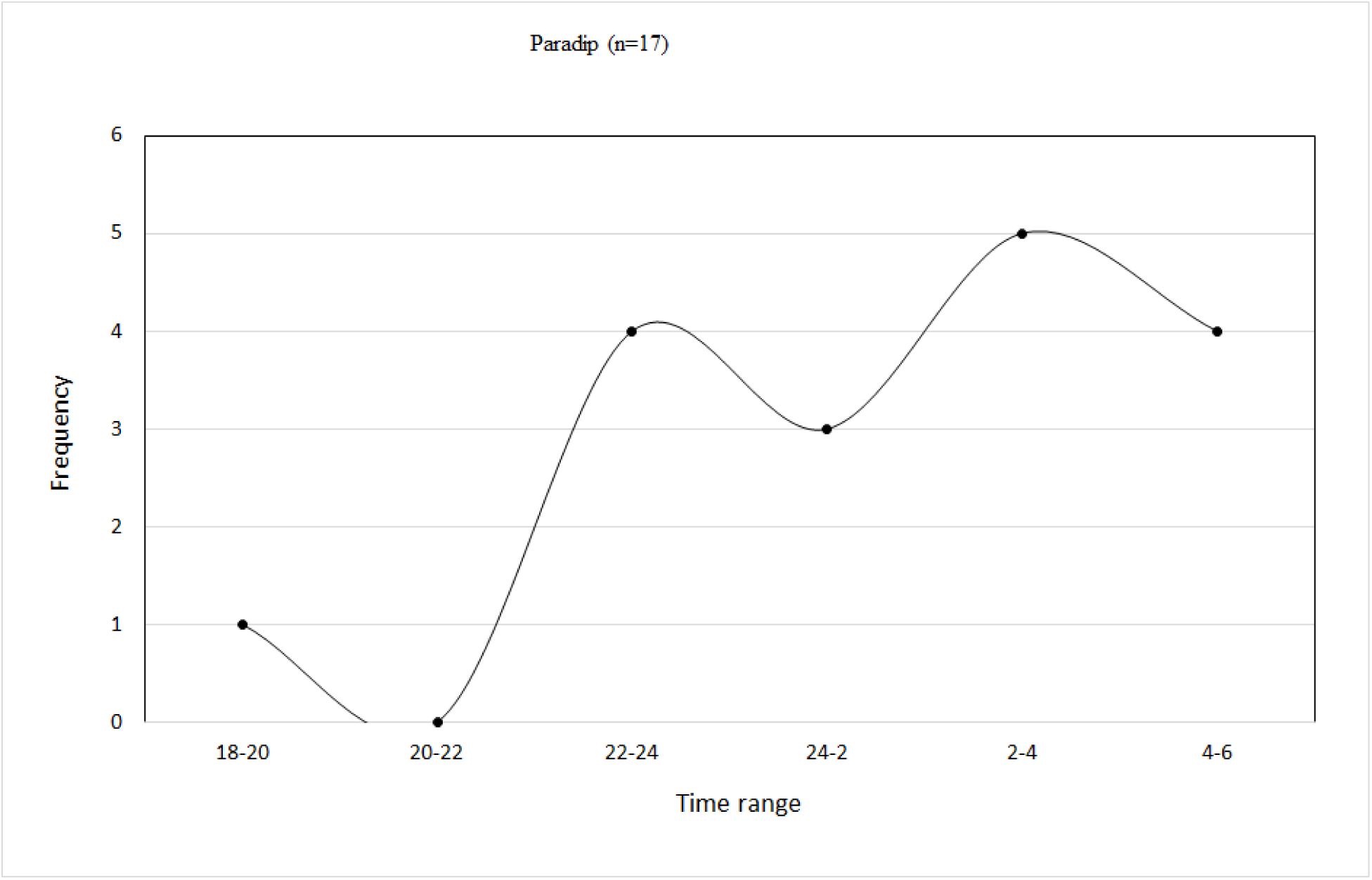
Graph showing activity pattern of Fishing Cat in Paradip

**Fig. 4c.**
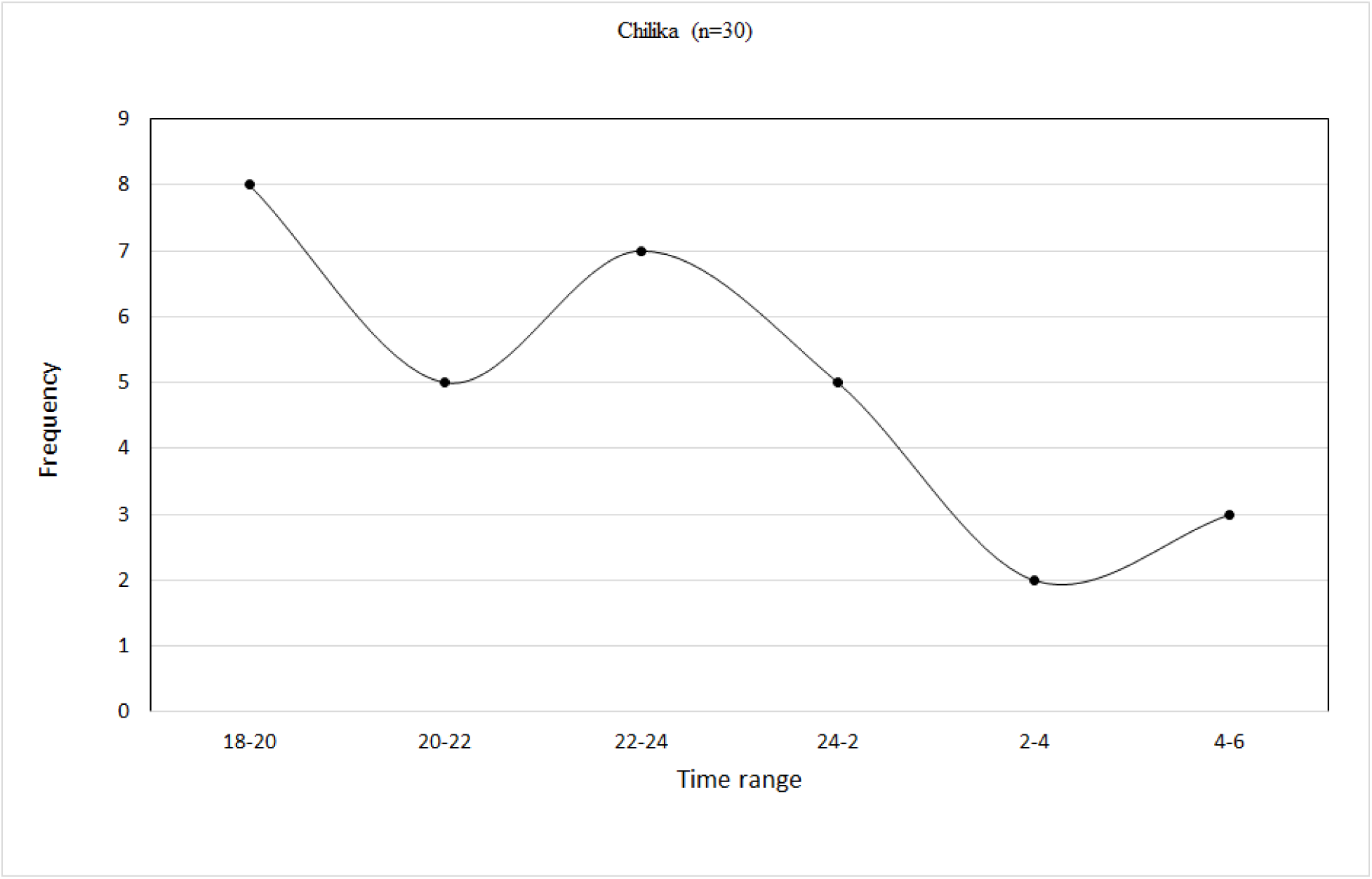
Graph showing activity pattern of Fishing Cat in Chilika

**Fig. 4d.**
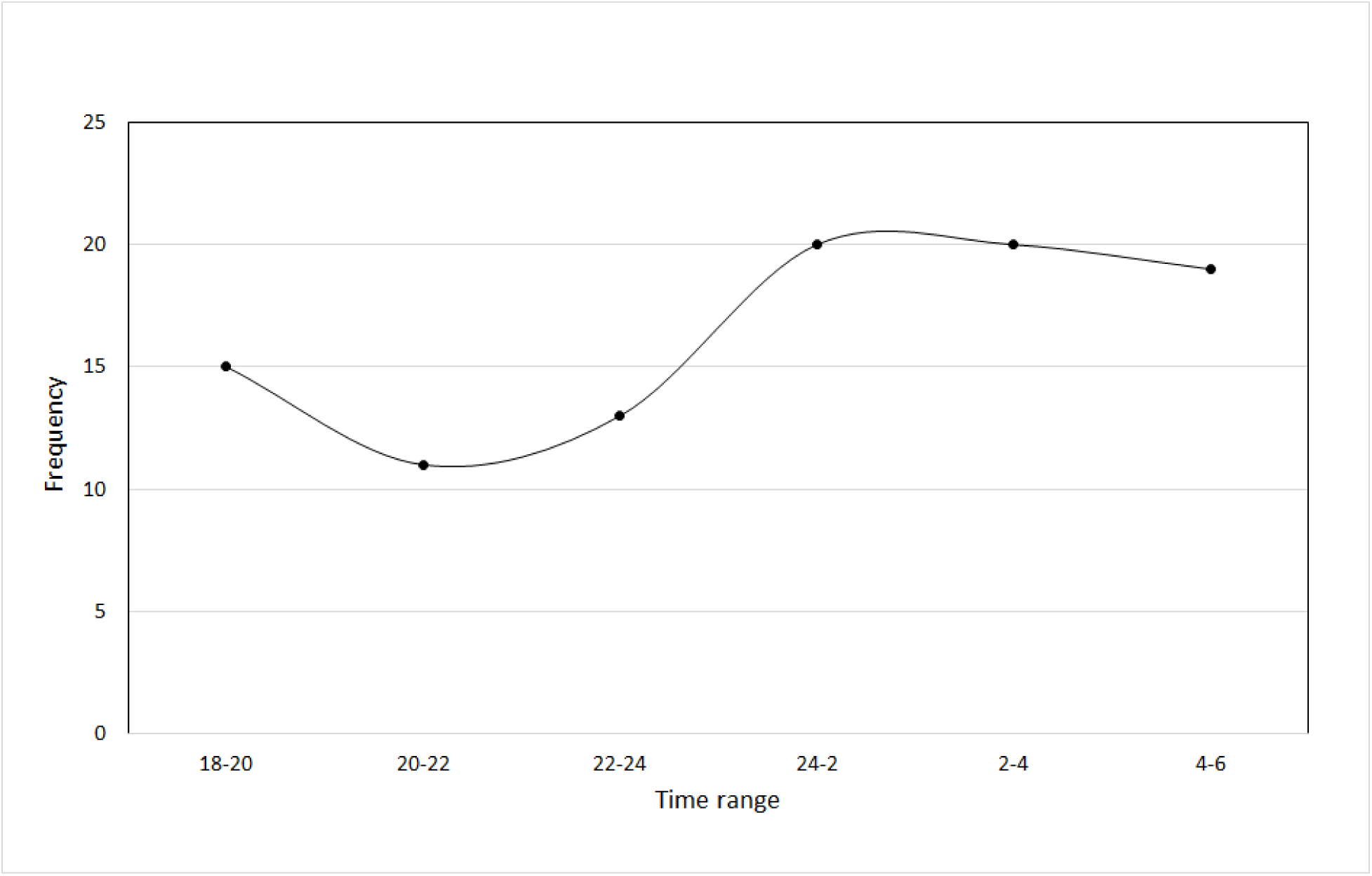
Graph showing general activity pattern (Howrah, Paradip and Chilika combined) of Fishing Cat outside Protected Area (n=98)

## Discussions

Most of Fishing Cat’s foraging time was spent in searching for prey (89.86%) in and around resource patches. While searching for prey in deeper waters which required submergence of the upper body, the cat alternated between mobile (37.94%) and stationary searching modes (51.65%). When no prey was detected the cat shifted to mobile searching again. In the event of an unsuccessful hunt, Fishing Cat came out of the waterbody, shook off the excess water and took position immediately at a different spot. Sometimes it also returned to the same spot. These findings are similar to an anecdote from Keoladeo Ghana Bird Sanctuary where it was observed to wait along the water’s edge concentrating for signs of prey and was found to change positions roughly every 15 minutes (Sunquist and Sunquist, 2017). In contrast, our study shows that a different strategy in shallow waters that did not require the upper portion of the body to get wet. Here, it deployed an active mode of searching for prey, occasionally pawing the waters to flush them out. This suggests that the cat switches from a combination of stationary- and-mobile mode of searching for prey in deeper waters to a more mobile and active mode of hunting in shallow waters in response to the differing needs to conserve energy.

Contrary to terrestrial foragers, aquatic foragers face additional physiological constraints (Rosen et al. 2007). For example, semi-aquatic foragers such as otter, muskrat, platypus lose body heat while foraging in water (Grant and Dawson 1978;Sherer and Wunder 1979;MacArthur 1984;Carss 1995;Kruuk 2006). Further, predator avoidance strategies of aquatic prey like fish (Katzir and Camhi 1993;Harvey and Nakamoto 2013) must be accounted for during a successfull hunt. Hence, a successful hunting strategy must compensate for energy loss while optimizing energy gain. For example, our data shows that the cat took measured attempts to hunt in deeper waters, i.e., plunge/dive into water with outstretched forelimbs to catch prey. The ‘attempt to hunt’ in deeper waters was found to be as low as 2.75%. This might be a strategy to maximize chances of prey capture while minimizing energy loss since each attempt to hunt leads to energy loss and too many attempts may alert the prey. The cat was found to be successful 60% of the time it attempted to hunt in deeper waters. We did not record a successful hunt from shallow waters but do not imply that Fishing Cat hunts more successfully in deeper waters. The findings are rather an artefact of the low-sample size of data from shallow water hunting. Irrespective of this, the chances of offsetting the high cost associated with attempting to hunt is variable and thus strategies to conserve energy must evolve.

Felids are known to mitigate high hunting costs by reducing energy loss in other stages of hunting, such as during pre-kill stages (Williams et al. 2014). The pre-kill stages constitute efforts to detect prey by searching and then approaching/stalking to reduce distance to prey after detection. While the ‘sit-and-wait’ search approach conserves energy, moderate energy might be spent in actively searching for prey. Upon detecting prey, stalking is used selectively to reduce distance to prey and increase chances of a successful hunt and in itself causes low energy loss. The kill stage (attempt to hunt) incurs a high cost in deeper water as against a relatively moderate cost in shallow water. Post-kill stages of fish consumption include subduing prey, processing, consuming and digesting. After subduing prey, some energy is spent in dragging a big fish (1 – 2 kg) as was interpreted from a field observation by one of the authors and from online repository (Bk Jeong 2017). Smaller prey may be consumed immediately next to the waterbody as was observed in camera traps. However, fish once caught are easy to subdue as against terrestrial prey (Sunquist and Sunquist 2017). Additionally, there is minimal cost of processing hunted fish since scales are mostly excreted out along with scats. More importantly, fish protein is easily digestible despite having high biological value (Vladau et al. 2008;Yilmaz et al. 2018). Thus, in deeper waters, the high cost of hunting is offset by adopting energy-conserving pre-kill stages, economic attempts to hunt and finally by successfully hunting highly-nutritious feed. In shallow waters, more time is spent in actively searching for food which would probably increase the chances of finding one and once a fish is successfully hunted, it could outweigh the energy spent in actively searching for food. This perhaps serves to answer why Fishing Cat prefers fish over rodents, which provide maximum metabolizable energy to small felids in general (Haque and Vijayan 1993; Mukherjee et al. 2004; Hunter 2019). Thus, while the ‘sit-and-wait’ (Supplementary Data SD 7) and ‘active foraging’ strategy (Supplementary Data SD 9) from the small cat lineage forms the backbone of Fishing Cat’s hunting strategy, its morphological adaptations to hunt prey in water further enhances the success of this strategy. Such adaptations are common among members of the carnivore guild that exhibit convergent evolution to forage in the ‘semi-aquatic niche’ (Hunter 2019).

In addition, we recorded scent marking and faecal deposition, which are known territory marking behaviors in felids (Brown and Bradshaw 2014;Burgos et al. 2018). We also recorded sniffing behavior near resource patches. This is interesting as the strength of the odor in male cat pheromone communicates the presence of high quality resource patches which is an attractant for potential mates (Brown and Bradshaw 2014) whereas sniffing helps in sensing the presence of other individuals as well as in gathering information on food resources (Geertsema 1984;B Bradshaw 2014). However, multiple individuals were also recorded to use the same site same night in three instances. This could be because of the relatedness of these individuals. While captive studies indicate possible male parental care (Rijsdijk 2011;Castaneda, [Cincinnati Zoo and Botanical Garden, Cincinnati, United States of America], personal communication, [December, 2019]) our data also shows sub-adults hunting in the presence of the mother. Thus, despite being solitary hunters, related individuals could be using the same resource patch, probably because of high density of prey, which is apparent from the use of such wetlands for fishing/pisciculture by humans.

The strictly nocturnal activity of fishing cat in our study area could be attributed to two reasons: a) the shift in activity pattern with intensifying levels of human presence could have to do with avoidance of humans, much like how it adjusts to daytime hunting in order to co-exist with larger competitors like Tiger and Leopard (Nair 2012) b) its crepuscular and nocturnal activity could also be attributed to activity spikes in fish, its primary prey (Carss 1995; Banerjee 2018).

The major caveat in our study is the post-hoc analysis of opportunistically collected data as opposed to data derived from a structured design with predefined objectives. It is unlikely that the major conclusions of the study will vary, nonetheless, more sampling from various seasons and landscapes will reveal finer details which could have ecological and conservation implications, the latter is particularly relevant in human dominated landscapes. For example, we would expect variations in ‘attempts to hunt’ during cold and warm seasons because heat loss might be less of a challenge in the latter. Locals also report greater fish loss during summer months when water levels in ponds decrease and hypoxic conditions prevail. Hence, quantifying ‘attempts to hunt’ and ‘successful hunts’ across seasons could help focus management interventions to minimize negative interactions between fish cultivators and Fishing Cat.

Finally, prey is central to predator ecology. Analyzing aspects of fish behavior with respect to hunting by Fishing Cat will thus add a vital dimension. For example, anecdotal accounts suggest that the cat taps the surface of the water to attract fish. Moreover, adult Fishing Cat use their paws to feel for fish in muddy waters and in a captive facility, a blind Fishing Cat was seen to feel around for food with his feet before plunging his head into the water to retrieve the food (Buck [Aspinall Foundation, Lympne, United Kingdom], personal communication, [July, 2011])). Additionally, hydrophytic vegetation provides refuge and congregation sites for fish (Romare and Hansson 2003;Sohel and Lindstrom 2015). Predatory fishes frequent hydrophytic vegetation in search of prey (O’Hara 2012;Ren et al. 2019). Therefore, the presence of such vegetation might be an important cue for hunting in both deeper and shallow waters. Future studies which consider avoidance strategies of fish could significantly enhance our knowledge on its hunting strategy.

Despite its caveats, the study is exemplary in demonstrating how by-catch data from participatory-science initiatives could reveal hitherto unknown aspects of behavioral ecology while also successfully circumventing the problems of equipment theft and need of greater manpower to work in human dominated landscapes. Furthermore, it reveals how Fishing Cat adapts to heavy human presence by shifting its foraging time to a chiefly nocturnal practice. The cat could well be taking advantage of human-used fishing gears like bamboo fish-baskets which have unidirectional valves through which fish enters and gets trapped as they were found to perch on beds of dried emergent vegetation formed on banks along which the baskets were deployed in ‘anticipation of prey’. Although we did not record Fishing Cat taking fish from the baskets, locals reported that both Fishing Cat and Otter take the entrapped fish. How piscivorous mammals take advantage of human-used gears requires further exploration as the strategy they deploy gives them access to easy food but also incites negative interactions with fishermen.

## Acknowledgement

The authors would like to acknowledge all participants of the ‘Know Thy Neighbours’ program - a participatory science initiative funded by Wildlife Trust of India, Wildlife Conservation Trust and Mohamed Bin Zayed Species Conservation Fund. We would like to extend our gratitude to student volunteers Ryan Rodrigues and Shreya Bhattacharya for initiating the camera trapping exercise and would especially like to thank Subrata Maity, Anil Maity, Bappa Malik, Maidul Islam Khan, Joydeb Pradhan, Sudhin Adhikari, Indrajit Adak, Saswat Pati, Sourav Pati, Kaloshi Behera, Baraju Behera and his team members from Ma Pakshi Suraksha Samity and members of Mahavir Pakshi Suraksha Samity whose enthusiastic participation is especially mention worthy. We would also like to thank Iravatee Majgaonkar, Partha Dey for their vital inputs in developing the manuscript. A special thanks to Dr. Kulbhushansingh Suryawanshi who asked a question, during a discussion over the preliminary findings of the study, which piqued us to re-look at our analysis and discover aspects that excited us.

## Supplementary Data

**Supplementary Data SD 1.**—A video showing Fishing Cat fast-walking along waterbody **Supplementary Data SD 2.** A video showing Fishing Cat slow-walking and searching along waterbody

**Supplementary Data SD 3.**—A video showing Fishing Cat switching between mobile (walking, approaching) and stationary (standing) searching modes along waterbody

**Supplementary Data SD 4.**—A video showing Fishing Cat sniffing and approaching towards edge of waterbody followed by employing stationary searching

**Supplementary Data SD 5.**—A video showing Fishing Cat approaching waterbody edge **Supplementary Data SD 6.** A video showing Fishing Cat employing mobile stalking along waterbody

**Supplementary Data SD 7.**—A video showing Fishing Cat employing stationary searching (sitting), followed by stationary stalking along waterbody, followed by attacking (misses the catch), pheromone spraying and taking position again at another spot

**Supplementary Data SD 8.**—A video showing Fishing Cat subduing and eating small fish along waterbody

**Supplementary Data SD 9.**—A video showing Fishing Cat employing active search in shallow water

**Supplementary Data SD 10.**—A video showing Fishing Cat engage in self-grooming along waterbody

**Supplementary Data SD 11.**—A video showing Fishing Cat swimming out of frame into waterbody

**Supplementary Data SD 12.**—A video showing Fishing Cat sniffing and depositing faeces along water canal

**Supplementary Data SD 13.**—A video showing Fishing Cat yawning along waterbody **Supplementary Data SD 14.** A video showing two Fishing Cats seemingly communicating along waterbody

**Supplementary Data SD 15.**—A video showing two sub-adult Fishing Cats employing active search in shallow water as the mother watches over from some distance

**Supplementary Data SD 16.**—A video showing a male and a female Fishing Cat walking along waterbody together

